# Impaired Spatiotemporal Encoding of Social Behavior and Anxiety in the Prefrontal Cortex of Mice Lacking ASD-Risk Gene *Shank3*

**DOI:** 10.1101/2025.07.15.664892

**Authors:** Hailee Walker, Damhyeon Kwak, Prakash Devaraju, Bethany C. Curd, Amaya M. Chikuni, Nicholas A. Frost

**Affiliations:** University of Utah, Department of Neurology, Salt Lake City, Utah, United States of America

**Keywords:** *SHANK3*, Autism, prefrontal cortex, ensemble encoding, social behavior, anxiety-related behavior, calcium imaging, network excitability

## Abstract

The prefrontal cortex is a central regulator of complex behaviors, including social interaction and anxiety-related behaviors. The prefrontal cortex encodes these behaviors using heterogeneous groups of neurons, or ensembles, which collectively process inputs and communicate with distributed brain regions. We examined whether loss of the Autism-risk gene Shank3 alters the recruitment of neurons encoding socioemotional behavior collectively, or if abnormal activity during specific behaviors might affect functionally or anatomically defined populations of neurons. To do this, we combined spatially-resolved microendoscopic calcium imaging across the prefrontal microcircuit with functionally defined labeling of neurons as control and mutant mice engaged in social interaction or anxiety-provoking behaviors. We then utilized a non-biased transcriptomic method to identify neurons activated by social interactions. We show that the recruitment of heterogeneous neuronal populations are altered in a cell type and spatially dependent manner by loss of Shank3, with impaired recruitment of behavior-specific activity patterns within superficial, but not deeper aspects of the prefrontal cortex.

## Introduction

Autism Spectrum Disorders (ASD) impact 1/36 children^1^ and arise from heterogeneous environmental and genetic^2^ factors which disrupt normal social interactions and behavior. Although increasing numbers of genetic perturbations are linked to ASD, several genes, including *SHANK3*, account for a substantial number of cases^3^. Anxiety and other comorbid neuropsychiatric conditions are observed frequently^4^, potentially implicating overlapping circuitry. In particular, the medial prefrontal cortex (mPFC) plays critical roles in regulating both social interactions as well as anxiety and is implicated in behavioral manifestations of ASD. Underlining the role of the mPFC in regulating social interactions, modulation of mPFC circuit activity may impair^5^ or rescue^6^ normal social behavior.

These findings imply that the mPFC performs specific computations or types of computations critical to processing information relevant to social behavior. Disruption of these computations may link genetic perturbations associated with ASD to abnormal behavior. Indeed, imbalanced excitation and inhibition (E/I imbalance) is observed across human patients and rodent models of ASD^7,8^, which disrupts the normal functioning of cortical circuits^9^ underlying social behavior^5,10,11^ and at its most extreme is associated with comorbid epilepsy in patients with ASD^12,13^.

*SHANK3* encodes a postsynaptic scaffolding protein that is disrupted in Phelan-McDermid Syndrome and is associated with ASD and intellectual disability. Intracortical recordings from mice modeling loss of Shank3 reveal abnormally increased activity in the mPFC during social interaction^9^. This abnormal activity was associated with abnormal ensemble recruitment during social behavior, suggesting that neurons were recruited inappropriately during behavior. Thus, abnormal excitability of prefrontal circuits lacking Shank3 drives elevated activity during social interaction. However, it is not known if this abnormal activity affects all neurons within the circuit equally, or if specific subpopulations might be affected heterogeneously. This distinction is potentially important because different types of neurons in different layers of the mPFC microcircuit have distinct functional properties^14^, connectivity^15^, transcriptional machinery^16^, and projection targets^17^. In other words, different types of neurons in different layers of the cortex play distinct functional roles in local and distributed circuit function.

We hypothesized that abnormal ensemble recruitment in mice lacking Shank3 might disrupt the generation of distinct neuronal ensembles representing different types of information, or the precision with which specific types of neurons or projections are recruited during different types of behavior.

To examine this question, we utilized *in vivo* calcium imaging of functionally labeled neurons within different anatomical locations within the mPFC in freely moving mice engaged in social interactions and anxiety-related behaviors. As in our previous work^18^ we show that ensembles recruited during social interaction and anxiety-related behaviors were distinct from each other and were differentially distributed across the extent of the mPFC. Notably, while prefrontal activity was increased only during social interaction, the spatial distribution of both ensembles was altered in Shank3 mutant mice by excessive recruitment biased toward the more superficial aspect of the mPFC. We observed similar changes in activity in Shank3 mutant mice using a non-biased transcriptional approach.

This abnormal recruitment of superficial neurons was associated with a loss of the specific recruitment of projections from the superficial aspect of the mPFC, resulting in increased overlap of activity during social interaction and anxiety-related behaviors.

These findings highlight the fact that E/I imbalance in mice lacking Shank3 affects populations of neurons heterogeneously. Furthermore, identifying specific populations of neurons that are abnormally excited may be critical to understanding how circuit dysfunction affects information processing underlying complex behaviors in these disorders.

## Results

We have previously observed increased activity in the mPFC of Shank3 knockout mice (KO) compared to their wildtype (WT) littermates^9^. We reasoned that this increased recruitment might be a general phenomenon related to abnormal excitability of the prefrontal circuit as a whole, or that abnormal activity might be restricted to specific components of the microcircuit. Similarly, circuit architecture might be altered in a manner that alters activity during social behavior, without affecting activity during other behaviors.

To examine these possibilities, we expressed GCaMP7f pan-neuronally under control of the Syn promoter and implanted microendoscopes into the mPFC of WT and KO mice (**Figure 1A-B**). This permitted us to record and quantify the activity of individual neurons and groups of neurons as the mice engaged in freely moving social interaction and anxiety-related behaviors. Recordings were performed throughout a paradigm where mice were alone in their home cage for 10 minutes before and after a 10-minute social session in which a novel sex-matched conspecific was introduced to the home cage. Following the second home cage epoch, mice were transferred to an Elevated Zero Maze (EZM) and allowed to freely explore the maze for thirty minutes. The EZM is a raised circular platform with alternating closed and anxiety-provoking open regions used to model anxiety-related behaviors. Together, this experimental paradigm permitted quantification of activity during home cage epochs, periods of direct social interaction (Social), periods in which the conspecific was present, but the mice were not interacting (non-Social), and periods of open and closed arm exploration within the EZM.

**Figure 1.**
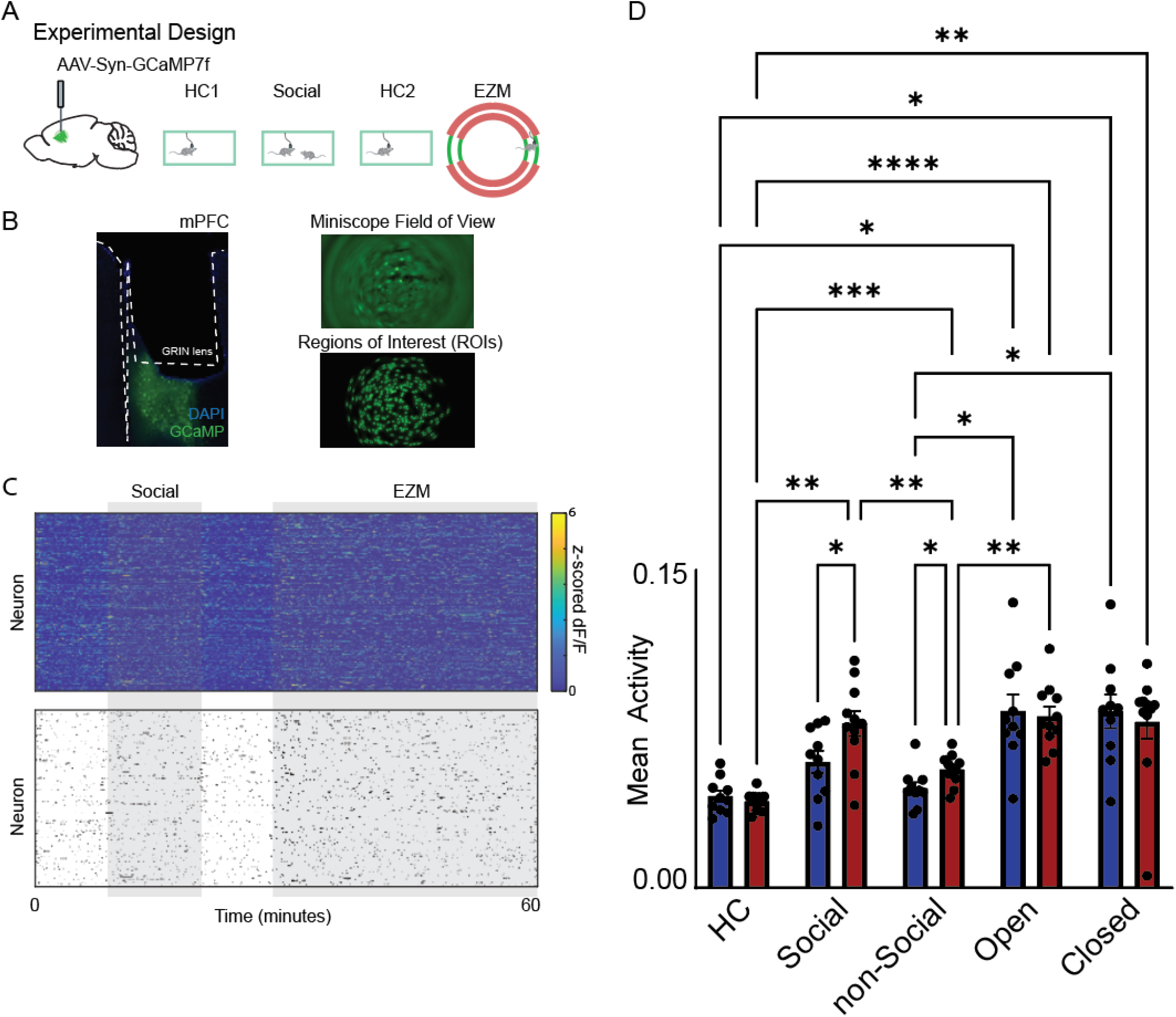
Abnormal mPFC ensemble recruitment during social interaction in Shank3 KO mice. (A) To monitor neuronal activity in the medial prefrontal cortex (mPFC) during social interaction and anxiety-related behavior, we injected AAV1-hSyn-GCaMP7f into the mPFC and implanted GRIN lenses into the ipsilateral mPFC. Mice were then recorded during social interaction followed by exploration of the elevated zero maze (EZM). (B) Left: representative histological section showing GRIN lens placement and GCaMP expression. Top right: representative field of view (FOV); bottom right: regions of interest (ROIs) extracted using CNMF-E. (C) Example calcium traces were binarized into raster plots representing active and inactive states for each neuron over time. (D) Bar graph showing the mean neuronal activity per mouse during social interaction and EZM exploration. Data include *n* = 1,843 neurons from 11 WT mice and 1,390 neurons from 11 KO mice (*n* reflects individual mice).

In total, we recorded activity from 1843 cells from 10 WT mice, and 1390 cells from 11 KO mice (mean 184.3 +/35.4 WT cells per mouse and 126.4 +/−17.8 KO cells per mouse; (**Supplementary Figure 1)**. As expected, both WT and KO mice spent a substantial amount of time during the social epoch engaged in social behavior. There was a trend toward less overall interaction time both at 10 minutes (301.6 +/−29.7 s vs 269.1 +/−33.0 s, p = 0.5, *t*-test, n = 10 WT mice and n = 11 KO mice) and in the initial 5 minutes (199.7+/−19.1 s vs 165.9+/−20.6 s, = = 0.2, *t*-test, n = 10 WT and n = 11 KO mice). WT and KO mice explored both the open and closed arms of the EZM equally (47.7 +/−3.8% WT vs 52.5+/−2.8% KO, p = 0.32, *t*-test, n = 10 WT mice and n = 11 KO mice; **Supplementary Figure 2**). Individual neurons were identified using CNMF-E and converted to a binary raster corresponding to periods in which each neuron was active or inactive (**Figure 1B-C**). We next quantified the percentage of time in which each neuron was active by calculating the proportion of frames in which the neuron was active and then averaging across mice. This permitted us to compare activity in WT and KO neurons across different epochs.

In KO mice, activity was similar to WT mice during HC epochs and increased significantly during EZM exploration to levels similar to those observed in WT mice. Activity significantly increased in both arms of the EZM. During social behavior, activity increased to levels similar to that observed during EZM exploration. In WT mice, we observed a trend toward increased mean activity during social interaction which was significantly increased in KO mice (**Figure 1D**). We also observed a significant increase in activity during periods in which the conspecific was present but the mice were not directly interacting (non-Social). Thus, the activity of prefrontal neurons in KO mice is specifically elevated (compared to WT mice) during social interaction.

We reasoned that this abnormal activity might be universally distributed within different layers of the mPFC, or might disproportionately affect different subpopulations of neurons within the laminar cortex. To investigate this question, we considered the activity of each neuron with respect to its location within the optical recording. The geometry of the mPFC is such that the implanted GRIN lens is situated orthogonally to the cortical microcircuit, allowing the simultaneous imaging of neurons located along the entire extent of the cortical microcircuit from superficial to deeper layers (**Figure 2A-B**). Thus, each of our recordings contains neurons located in both superficial and deeper aspects of the mPFC, permitting us to simultaneously quantify the activity of neurons localized across the laminated mPFC microcircuit.

**Figure 2.**
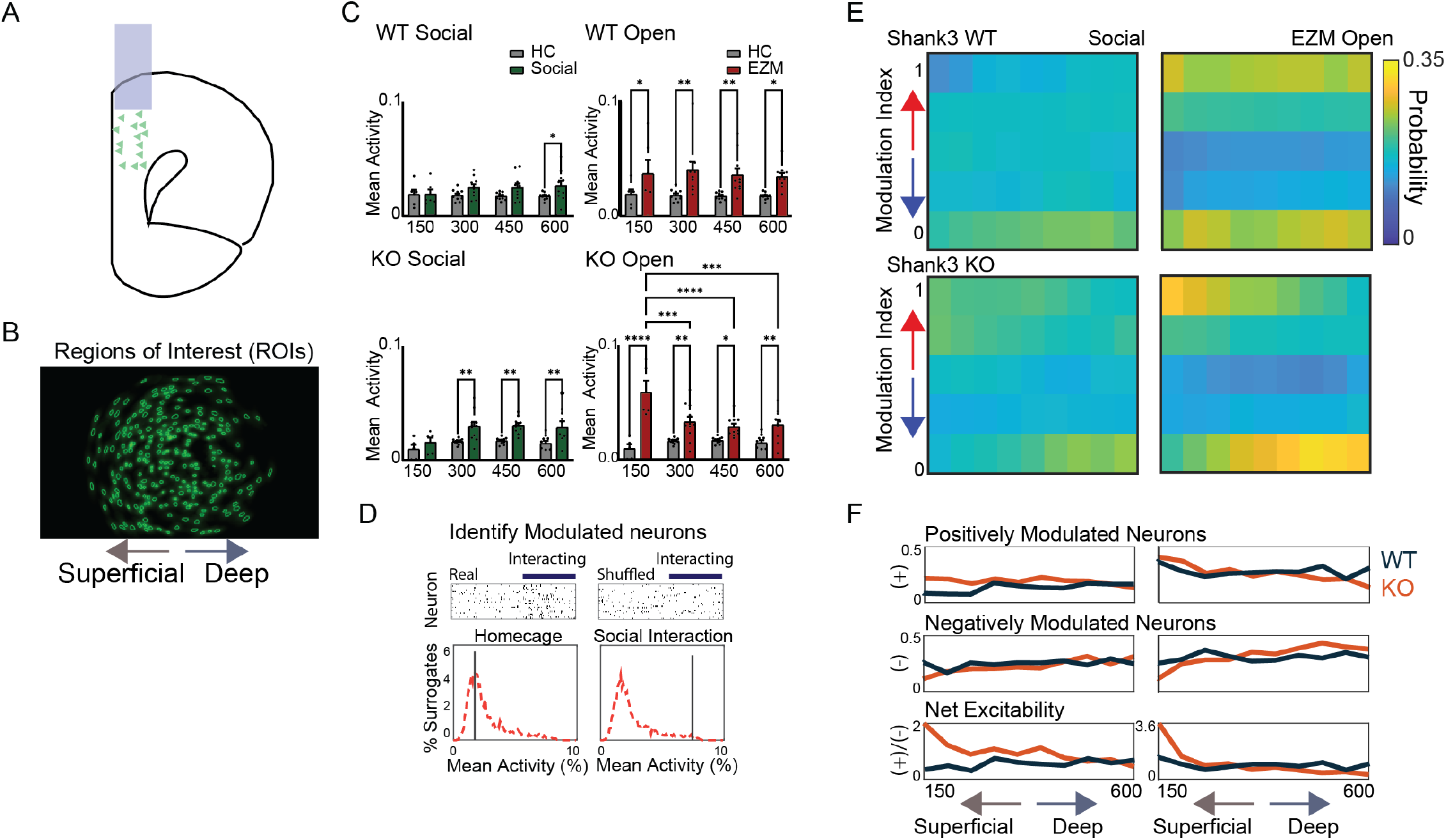
Distinct spatial profiles of social and anxiety-related ensembles are disrupted in Shank3 KO mice. (A) Schematic illustrating the placement of the GRIN lens relative to the cytoarchitecture of the medial prefrontal cortex (mPFC). (B) Representative field of view showing regions of interest (ROIs) from a single microendoscopic calcium imaging session. The left edge of the field corresponds to the medial/superficial aspect of the mPFC. (C) Mean neuronal activity during social interaction and anxiety-related behavior (elevated zero maze), plotted as a function of distance from the superficial edge of the GRIN lens, binned in 150 μm increments. In both WT and KO mice, superficial neurons showed increased activity during anxiety; however, only KO mice showed increased superficial activity during social interaction. (D) Behavioral modulation was assessed for each neuron by comparing its activity to a null distribution generated from temporally shuffled data (red). (E) Proportion of neurons that were positively (top row) or negatively (bottom row) modulated by behavior, plotted by depth from superficial to deep mPFC. (F) Quantification of the number of positively or negatively modulated neurons, and the positive-to-negative modulation ratio across depth bins. Data include *n* = 1,843 neurons from 10 WT mice and 1,390 neurons from 11 KO mice.

Our approach was to first bin identified neurons into 150-micron bins across the lateral extent of the optical recording. We then calculated the mean activity of all neurons recorded within each bin during social and nonsocial epochs, as well as during exploration of the EZM open arm. As before, we observed a modest increase in the mean activity of WT neurons during social interaction epochs; this was only significantly elevated in the deepest aspect of the mPFC. In contrast, we observed a significant increase in the mean activity within all spatial bins of the mPFC during EZM exploration (**Figure 2C**), suggesting that exploration of the EZM open arm recruits distinct mPFC microcircuits compared to social interaction. Quantification of ensemble activity during social interaction and EZM exploration resulted in qualitatively similar results in KO mice. However, in KO mice, we observed significant elevations in the mean activity during social interaction within more superficially oriented spatial bins, and there was also increased activity in the most superficial spatial bin during EZM exploration (**Figure 2C**). Together, these findings suggested that loss of Shank3 increased cortical activity during both social interaction and EZM exploration in more superficially localized neurons.

We reasoned that changes in the mean activity reflect both neurons that are positively modulated, as well as neurons that are negatively modulated. As we have previously observed that the number of positively modulated neurons during social interaction is increased in KO mice, with a corresponding decrease in negatively modulated neurons^9^, we examined whether these changes in ensemble recruitment were restricted to specific aspects of the cortical microcircuit.

To accomplish this, we utilized our previous approach^9^ to identify neurons that were positively or negatively modulated during social interaction or EZM exploration by quantifying the activity of each neuron during each respective behavior and then comparing its real activity to a shuffled distribution. Neurons that are strongly positively modulated have high activity relative to their shuffled distribution (>90^th^ %ile), whereas negatively modulated neurons have relatively low activity compared to shuffled data (< 10^th^ %ile) (**Figure 2D**). By binning neurons into 100-micron spatial bins and then quantifying the proportion of neurons which were positively or negatively modulated, we were able to visualize spatially distinct patterns of ensemble recruitment composed of positively modulated and reciprocally inhibited neurons for each behavior across the extent of the mPFC (**Figure 2E**).

This approach again revealed spatially distinct patterns of ensemble recruitment across the extent of the mPFC during social interaction and EZM exploration. Similar to the mean activity, we observed increasing numbers of positively modulated neurons within deeper spatial bins of the mPFC during social interaction. However, this was coupled with increasing numbers of negatively modulated neurons in deeper aspects of the mPFC. This was distinct from the pattern observed during EZM exploration in which the proportion of positively and negatively modulated neurons was increased compared to social interaction across the extent of the mPFC.

Surprisingly, ensemble recruitment during both social interaction and EZM exploration followed a distinct spatial pattern in KO animals with increasing proportion of positively modulated neurons in superficial aspects of the mPFC. This was paired with a decrease in the proportion of negatively modulated neurons, resulting in increased net excitability in superficial aspects of the mPFC during both social interaction and EZM exploration (**Figure 2F**). Thus, ensemble recruitment during social interaction and EZM exploration is disrupted in the superficial aspect of the mPFC in mice lacking Shank3.

We further investigated cell-type specific changes in ensemble recruitment using a non-biased transcriptional approach which we have previously used to quantify the contribution of heterogeneous cell types to social encoding^19,20^. To do this, we isolated single nuclei from mPFC of WT and Shank3 KO mice following social interaction with 5 novel juveniles in the home cage, and generated cDNA libraries for 10X Genomics snRNA-sequencing. In total, we isolated and clustered 78,484 cells from 11 WT mice and 38,307 cells from 5 KO mice. These cells were clustered using Seurat and 20 cell types were identified using known marker genes (**Supplementary Figure 3A & E-G, Figure 3B**). All clusters were populated with a combination of cells from WT and KO mice (**Supplementary Figure 3B**). This included 7 excitatory neuron types (**Supplementary Figure 3C**), 5 inhibitory neuron types (**Supplementary Figure 3D**), and 8 non-neuronal. The excitatory neuron types were then grouped by layer (**Figure 3A**) to understand how the recruitment of neurons changes across layers following social interaction for each genotype. Interestingly, following clustering we observed quantitatively more layer 2/3 excitatory neurons in KO mice and fewer glial cells (**Figure 3C**). Other neuronal clusters were similar in quantity in WT and KO mice (**Figure 3C, Supplementary Figure 3H-I**).

**Figure 3:**
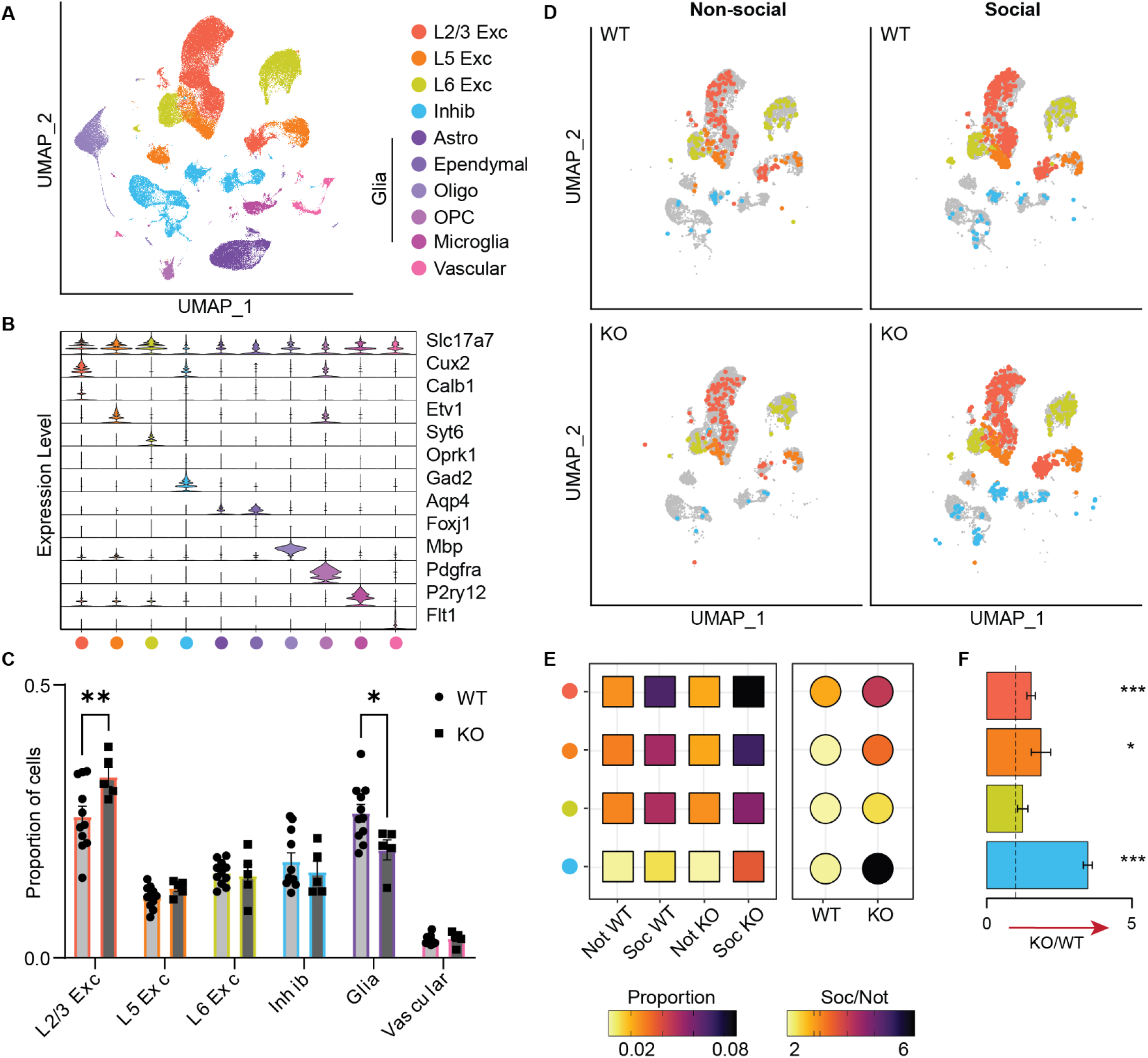
Social interaction increases ensemble recruitment in Shank3 Knock-outs as measured by immediate early gene expression. (A) UMAP of identified cell types (n = 116,791 cells; 11 WT and 5 KO mice). (B) Gene marker expression for each cell type. (C) Proportion of cells belonging to each cell type for WT (light grey) and KO (dark grey) mice (mean +/−SEM). (D) IEG+ neurons in control or social conditions demonstrate an increase in IEG+ neurons in both WT and KO mice. IEG+ cells are colored by cell type and overlaid on corresponding clusters from (A). Each experimental group was randomly down sampled to consist of the same number of cells per experimental group. (E) Boxes on the left are colored by the proportion of IEG+ cells for each genotype and cell type before and after social interaction. Circles show the ratio of IEG+ cells in the social condition over the baseline level for each genotype with darker colors showing a greater increase in proportion of active cells. (F) Magnitude of IEG positivity in Shank3 KO animals as compared to WT controls. Bar graph of the ratio of the fold-difference in the proportion of IEG+ neurons in each cell population plotted as WT/KO (Left side) or KO/WT (Right side) to show the increased IEG+ proportion in either KO or WT (L2/3 Exc KO/WT proportion 1.87 +/−0.33, p=0.001; L5 Exc KO/WT proportion 1.87 +/−0.02, p=0.02; L6 Exc KO/WT proportion 1.24 +/−0.18, p=0.1; Inhib KO/WT proportion 3.48 +/−0.15, p=0.001; P-values and standard deviations calculated by from a null distribution generated by shuffling real data 1,000 times; n = 55,183 WT neurons from 11 WT mice, 29,592 KO neurons from 5 KO mice).

Similar to our observations using implanted microendoscopes, a greater proportion of excitatory neurons in layer 2/3 (L2/3) and layer 5 (L5) are activated during social interaction in KO mice, whereas neurons located in layer 6 (L6) are activated in similar numbers in WT and KO mice (**Figure 3D-F**). As this approach allowed us to quantify the activity of a large number of transcriptionally-defined excitatory and inhibitory neuron populations, we next examined the degree to which these more specific neuronal populations were activated during social interactions. Superficial excitatory neuron populations consisted of L2/3, L5 and a population of excitatory markers expressing both L2/3 and L5 marker genes as identified by their expression of Cux2, Calb1, and Etv1^21^ Deeper layer excitatory neurons consisted of L6 Syt6+, L6 Oprk1+, L6b and L6 Car3+ cell types (**Supplementary Figure 3C & E**). Again, the three superficial excitatory neuron types showed increased recruitment during social interactions which occurs to a greater extent in the Shank3 KO mice (**Supplementary Figure 4A-B**).

Compared to excitatory neurons, inhibitory neuron populations were less likely to express IEGs before or after social interaction, as a group (**Figure 3D**). However, when we pooled inhibitory Gad2+ neurons we saw a dramatic increase in the proportion of IEG+ inhibitory neurons following social interaction in KO, but not WT mice (**Figure 3E-F**). We reasoned that the recruitment of inhibitory neurons might be globally increased, or the recruitment of specific subpopulations might be altered. Consistent with the latter hypothesis, PV and VIP interneurons seem to have increased recruitment in KO social ensembles, while somatostatin (Som) neurons showed reduced recruitment (**Supplementary Figure 4C-D**). This is consistent with prior studies showing Shank3-deficiency results in loss of inhibitory feedback from Somatostatin-positive neurons in prefrontal cortex and that Shank3 expression in inhibitory populations is critical for circuit function ^22,23^.

Together, these data suggest that abnormal ensemble recruitment in Shank3 KO mice is not evenly distributed within different anatomical locations and cell types. We reasoned that this might reflect broad changes in circuit function, or in fact alter the output of prefrontal circuits during behavior. Communication between the vHPC and mPFC is broadly important in regulating social behavior^24^, anxiety^25^, and spatial navigation^26^ and vHPC activity is altered during social interaction in Shank3 mutant mice^27^. Similarly, NAc projections from the mPFC are involved in encoding social interactions^17^ and cortico-striatal circuitry has been shown to be disrupted in Shank3-deficiency^28,29^. To examine these possibilities, we utilized a combinatorial genetic strategy to identify neurons monosynaptically innervated by inputs from the ventral hippocampus (vHPC), or neurons which projected to the nucleus accumbens (NAc) in subsets of WT and KO recordings.

Our labeling strategy was as follows: vHPC innervated neurons were labeled by injecting AAV1-hSyn-Cre into the vHPC of Shank3-mutants crossed with Ai75 mice such that a nuclear-localized tdTomato marker was expressed in neurons in the mPFC receiving monosynaptic innervation from AAV1-cre expressing vHPC projections. NAc projections were labeled utilizing retroAAV-hSyn-mCherry injected into the nucleus accumbens such that mPFC neurons projecting to this region were labeled with mCherry. AAV1-Syn-GCaMP7f was then injected in the mPFC and a GRIN lens was implanted 150 um above the injection site (**Figure 4A-B**). We then collected two color microendoscope images of GCaMP-expressing and static red-labeled neurons as mice engaged in social interaction and EZM exploration (**Figure 4C-D**). This generated a total of 1559 unlabeled neurons from 10 WT mice, 62 vHPC innervated neurons from 5 WT mice, and 51 NAc projecting neurons from 5 WT mice, as well as 1028 unlabeled neurons from 11 KO mice, 101 vHPC innervated neurons from 5 KO mice, and 45 NAc projecting neurons from 6 KO mice (155.9 +/−32.1 unlabeled cells/WT mouse vs 93.5 +/−13.8 unlabeled cells/KO mouse; 10.2+/−2.4 NAc labeled cells/WT mouse vs 7.5+/−2.4 NAc labeled cells/KO mouse; 12.4 +/−5.1 vHPC labeled cells/ WT mouse vs 20.2 +/−8.9 vHPC labeled cells/KO mouse) (**Supplementary Figure 2**).

**Figure 4.**
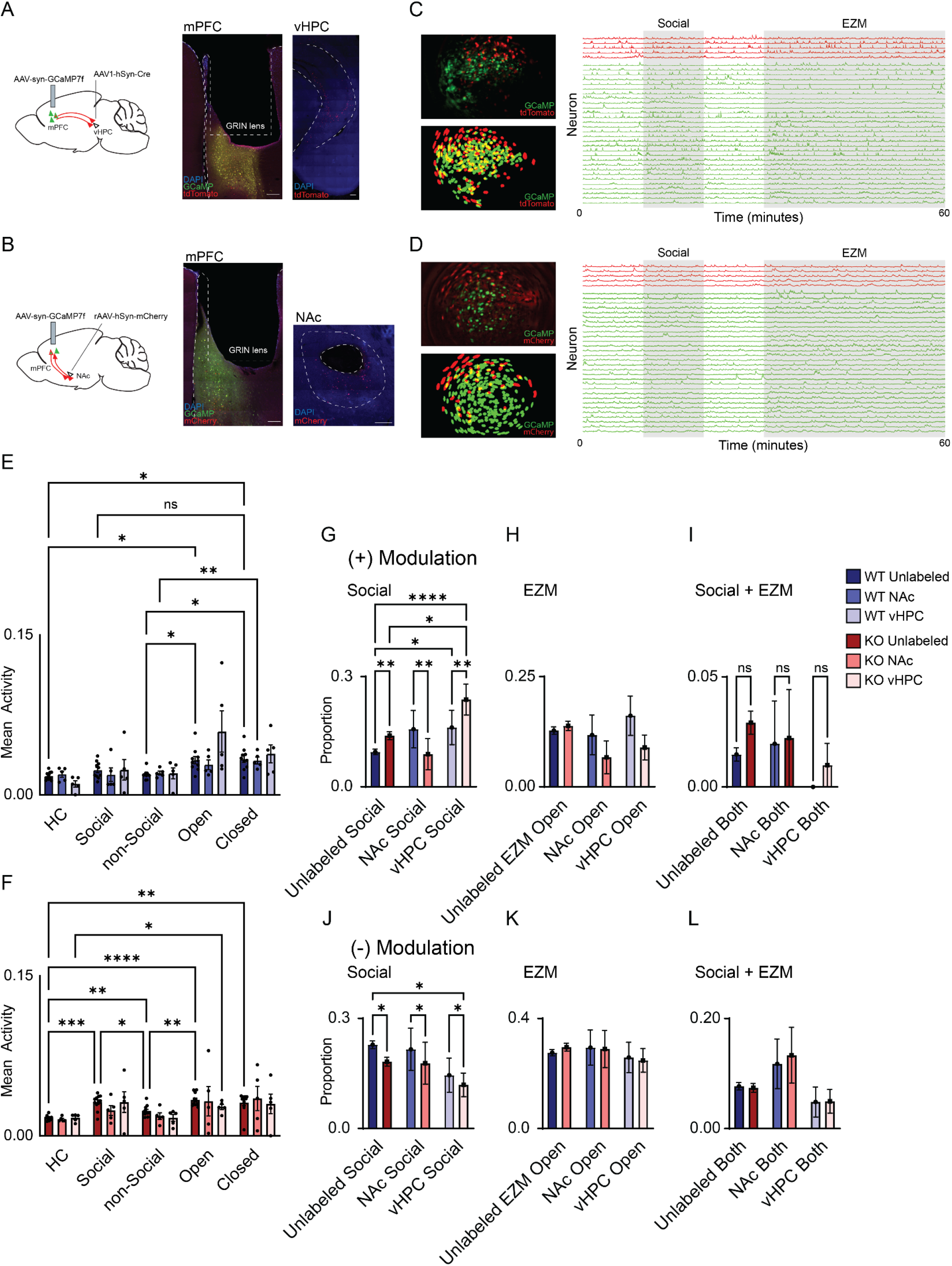
Abnormal recruitment of anatomically defined mPFC connections during social and anxietyrelated behaviors in Shank3 KO mice. We recorded the activity of anatomically defined mPFC neurons that either received monosynaptic input from the ventral hippocampus (vHPC) or projected to the nucleus accumbens (NAc) during behavior. (A) To label mPFC neurons innervated by vHPC inputs, we crossed Ai75 reporter mice with Shank3B mutant mice, then injected AAV1-hSyn-Cre into the vHPC of WT or KO mice. Since AAV1 is transported anterogradely, this resulted in tdTomato expression in mPFC neurons that receive vHPC input. A subsequent injection of AAV1-hSyn-GCaMP7f into the mPFC enabled expression of GCaMP7f in all mPFC neurons for imaging via an implanted GRIN lens. (B) To label mPFC neurons projecting to the NAc, we injected AAVretro-hSyn-mCherry into the NAc and AAV1-hSyn-GCaMP7f into the mPFC, allowing visualization of both projection identity and calcium dynamics. (C) Representative FOV of GCaMP transients (green) and static tdTomato (red) signals (top left), cell map with green ROIs extracted using CNMF-E and red ROIs generated manually to identify vHPC-innervated neurons (bottom left), and traces of neurons innervated by vHPC inputs (red) or not (green) (right). (D) Same as (C), but for static mCherry signals to identify NAc-projecting neurons. (E) Bar graph showing the mean activity of unlabeled, NAc-projecting, and vHPC-innervated mPFC neurons in WT mice. (F) Same as (E), but for Shank3 KO mice. (G) Bar graph showing the proportion of neurons in each group that were positively modulated during social interaction. (H) Bar graph showing the proportion of neurons positively modulated during exploration of the open arm of the elevated zero maze (EZM). (I) Bar graph showing the proportion of neurons in each group that were positively modulated during both social interaction and EZM open-arm exploration. (J-L) Same as (G-I), but for negatively modulated neurons. Data is from 1559 unlabeled neurons from 10 WT mice, 62 vHPC innervated neurons from 5 WT mice, and 51 NAc projecting neurons from 5 WT mice, 1028 unlabeled neurons from 11 KO mice, 101 vHPC innervated neurons from 5 KO mice, and 45 NAc projecting neurons from 6 KO mice.

This permitted us to quantify activity of vHPC innervated and NAc projecting neurons, in addition to unlabeled cells during social interaction and EZM exploration. As before we observed a greater increase in the mean activity of unlabeled neurons from KO mice (compared to WT controls) as well as labeled populations during social interaction, though this increase was not significant in either labeled population (**Figure 4E-F**). The activity of all three populations of neurons was right shifted in cumulative distribution plots drawn for KO cells compared to WT cells during social interaction (**Supplementary Figure 5**). This was driven by an increase in the proportion of unlabeled and vHPC-innervated neurons which were positively modulated as well as a decrease in the proportion of negatively modulated neurons in KO neurons from each subpopulation compared to their WT counterparts during social interaction. Notably, NAc projecting neurons from KO mice were less likely to be positively modulated during social interaction compared to their WT counterparts (**Figure 4G, J**), suggesting an impairment in circuit output. We observed no significant differences in the proportion of neurons which were positively or negatively modulated during EZM exploration (**Figure 4H, K**).

Using this definition of modulation (> 90% of shuffled, or < 10% of shuffled) we observed a small number of comodulated neurons (neurons which were positively modulated during both social interaction and EZM exploration (**Figure 4I, L**). We further explored this by quantifying the number of neurons which increased their activity in both social interaction and EZM exploration in superficial (< 300 microns) and deeper (> 300 microns) aspects of the mPFC microcircuit (**Figure 5**). Consistent with abnormal excitability (see Figure 2 and 3) in superficial aspects of the mPFC, we observed a greater number of KO neurons in superficial, but not deeper aspects of the mPFC were positively modulated by both social interaction and EZM exploration (**Figure 5C**). Thus, the precision with which ensembles can be generated for specific behaviors is impaired in the superficial aspect of the mPFC. Notably, while WT mice had similar numbers of co-modulated neurons in superficial and deeper aspects of the mPFC, unlabeled and NAc projecting neurons in superficial aspects of KO mPFC were more likely to be comodulated by the two behaviors (**Figure 5D**), suggesting that ensemble recruitment in the more superficial aspects of prefrontal cortex is specifically impaired.

**Figure 5.**
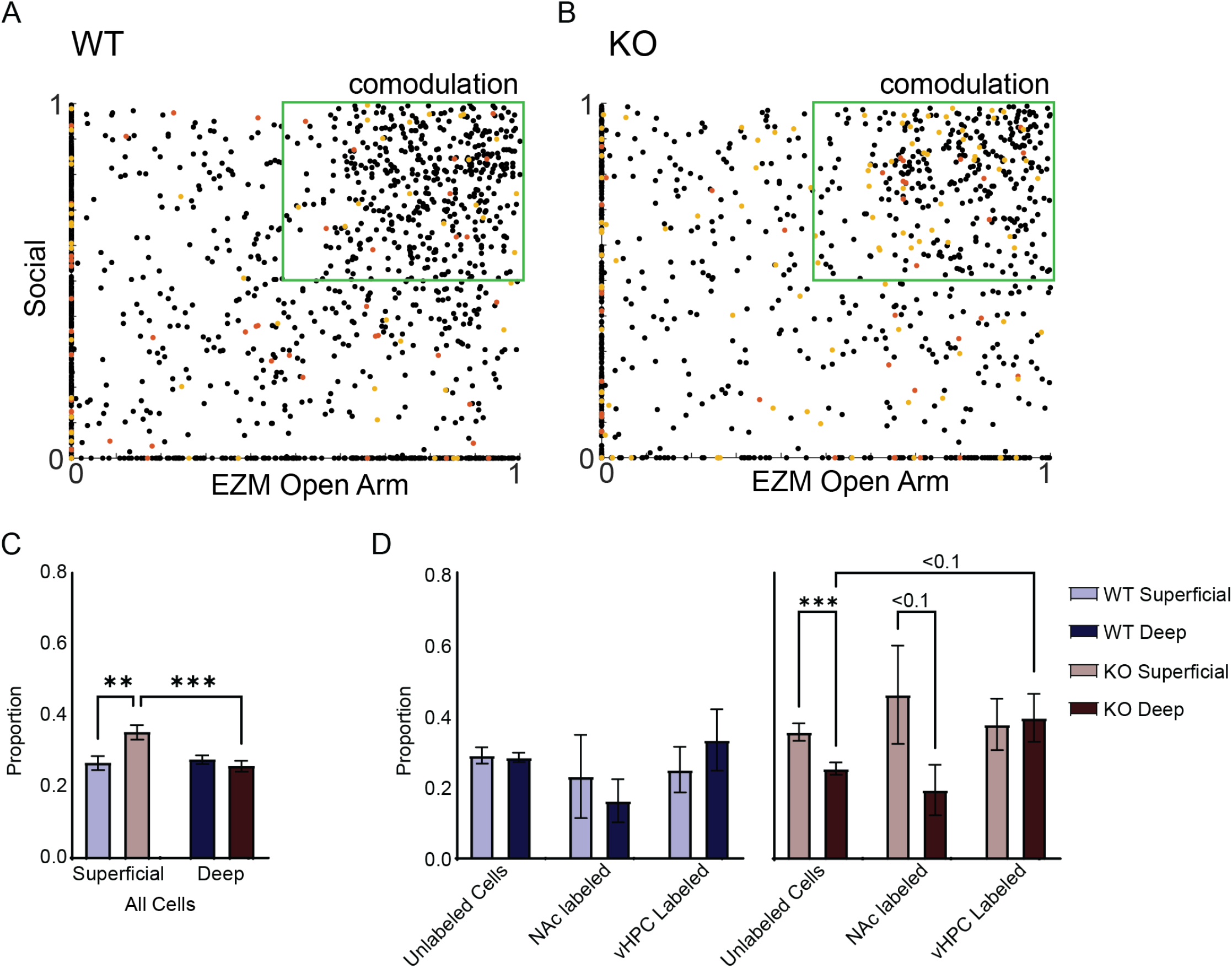
Loss of behavior specificity in superficial mPFC neurons of Shank3 KO mice. (A-B) Scatter plots showing the modulation index of individual neurons from WT and Shank3 KO mice during social interaction and elevated zero maze (EZM) exploration. Neurons were classified as co-modulated if they exhibited positive modulation (modulation index > 50th percentile) during both behaviors. (C) Quantification of the proportion of all neurons classified as co-modulated in WT and KO mice. Neurons located within 300 μm of the cortical surface were classified as superficial, and those deeper than 300 μm as deep. (D) Bar graphs showing the proportion of unlabeled, nucleus accumbens (NAc)-projecting, and ventral hippocampus (vHPC)-projecting neurons that were co-modulated by both behaviors. Data are segmented by cortical depth (superficial vs. deep) and genotype (WT vs. KO). Data is from 1559 unlabeled neurons from 10 WT mice, 62 vHPC innervated neurons from 5 WT mice, and 51 NAc projecting neurons from 5 WT mice, 1028 unlabeled neurons from 11 KO mice, 101 vHPC innervated neurons from 5 KO mice, and 45 NAc projecting neurons from 6 KO mice.

**Figure 6.**
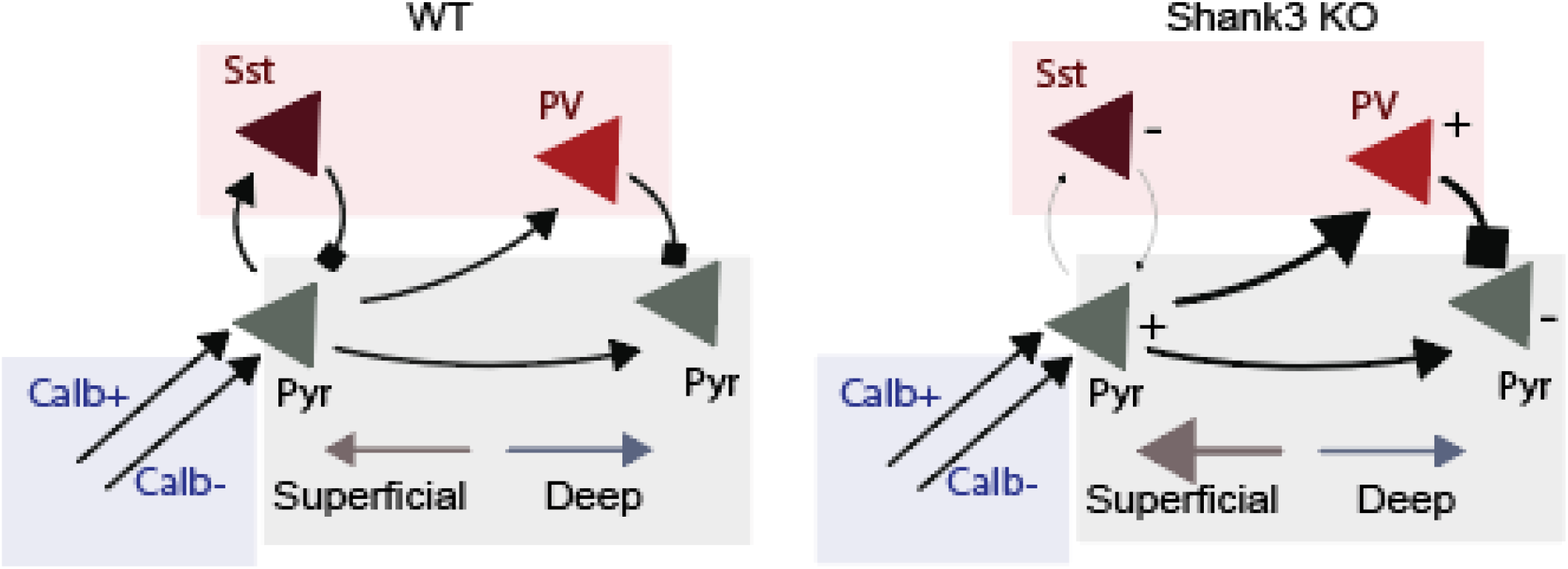
Model depicting how loss of Shank3 affects cortical dynamics. Schematic illustrating proposed circuit-level changes in medial prefrontal cortex (mPFC) associated with Shank3 deletion. In wild-type (WT) mice (left), superficial and deep pyramidal (Pyr) neurons exhibit behaviorally selective activation patterns. During social interaction, excitatory drive such as from calbindinpositive (Calb^+^) and calbindin-negative (Calb^−^) superficial inputs delivered from the vHPC is balanced by local inhibition from somatostatin-expressing (Sst) and parvalbumin-expressing (PV) interneurons. In Shank3 knockout (KO) mice (right), this balance is disrupted as superficial pyramidal neurons are hyperexcitable, leading to a loss of behavior-specific activation and driving increased feedforward inhibition onto deeper layer neurons. PV interneurons show increased influence (+), while Sst interneurons show reduced activity (−), leading to an altered inhibitory microcircuit.

## Discussion

Social interactions and anxiety-related behaviors are flexibly regulated by distributed cortical and subcortical regions including the mPFC. This flexibility requires integration of multiple types of information which may be encoded in parallel^18^ along with social information. Disrupted circuit function in ASD circuits^7-9,23,28,30,31^ may therefore impair computations required to generate distinct ensemble representations during different behaviors, ultimately driving pathologic changes in ASD-associated behaviors. Changes in Excitation/inhibition ratios have been well described in various ASD models, leading to increased excitability of cortical circuits^8,9,32^. However, it is not known whether changes in excitability affect circuits globally or in a cell-type dependent manner. This is an important question as heterogeneous neuronal populations in the mPFC have distinct functional properties^14^, connectivity^15^, transcriptional machinery^16^, and projection targets^17^ that may impact their role in the circuit and in behavior.

In this study, we investigate how the activity of neuronal populations defined by their location, gene expression, or anatomic connections is altered during social interactions and anxiety-related behaviors in the mPFC of mice lacking ASD-risk gene Shank3. We identified abnormally increased activity in the superficial aspect of the mPFC of Shank3 KO mice, during both social interaction and exploration of the open arm of the anxiety-provoking EZM. We further show that while activity is broadly increased across a variety of excitatory and inhibitory neurons, specific populations of excitatory neurons (particularly deeper-layer excitatory neurons, and Somatostatin-expressing interneurons) do not exhibit increased firing during these behaviors.

This suggests that microcircuits utilizing specific types of inhibition are not properly utilized in mice lacking Shank3 and is congruent with a previous report that Shank3-deficiency results in loss of inhibitory feedback from Somatostatin-positive neurons in prefrontal cortex ^23^ and that Shank3 expression in inhibitory populations is critical for circuit function^22^. Moreover, these results indicate that simply increasing inhibitory tone in a blanket fashion ^33^ may be insufficient to rescue behavioral deficits driven by selective loss of specific subpopulations of inhibitory neurons.

We next asked whether abnormal activity in the mPFC was internally generated or related to abnormal input from distributed brain regions (i.e. the vHPC). Projections from the vHPC to the mPFC have known roles in social behavior^24,34,35^, anxiety^25^, and spatial navigation^26^ and are specifically involved in dysregulated PV-interneuron recruitment during social interaction^27^. Additionally, previous work has demonstrated dysregulated vHPC function in Shank3 KO mice during social interactions. Highlighting the role of vHPC inputs in communicating social interaction, we observe significantly greater numbers of vHPC-innervated neurons are modulated during social interaction (compared to unlabeled neurons). Interestingly, we observed that mPFC neurons monosynaptically innervated by vHPC projections are more likely to be positively modulated during social interaction (compared to vHPC-innervated neurons in WT mice); however, these neurons are not disproportionately hyperactive (compared to the overall population).

Similarly, changes in mPFC neuronal activity likely impact downstream targets. The nucleus accumbens (NAc), which receives projections from the mPFC, has been implicated in the encoding of social interactions^17^. Prior work additionally shows that cortico-striatal circuitry is disrupted in Shank3-deficiency^28,29^ suggesting a potential role of this circuit in the altered social behavior observed in these mice. Our results indicate impaired mPFC output, evidenced by a reduction in the proportion of neurons modulated by social interaction, along with a loss of behavioral specificity in superficial mPFC layers. Notably, this impaired selectivity is present in both unlabeled and NAc-projecting neurons within the superficial—but not deep—layers of the mPFC. In contrast, we do not observe layer-specific differences in the behavioral selectivity of neurons innervated by the ventral hippocampus (vHPC).

While the design of our study enables investigation of layer- and cell type-specific contributions to altered mPFC activity, it does not permit dissection of the roles of specific subregions within the mPFC in driving behavioral differences. Our GRIN lens placement primarily targets the infralimbic portion of the mPFC, limiting our spatial resolution across subunits. Additionally, our transcriptomic method allows identification of neurons that are activated but not those that are inhibited. Although combining this approach with in vivo calcium imaging partially addresses this limitation, we remain unable to determine the specific contributions of individual cell types to the observed reduction in negatively modulated neurons.

Overall, our data suggests that there are specific computations that are impaired by loss of Shank3, and that understanding which populations of neurons at an anatomical and cell type level is important for fully understanding the effects of these and other ASD mutations on circuits. Moreover, individual behaviors such as anxiety and social behavior deficits may be caused by altered activity in differing neuronal populations, inputs, or targets rather than a single difference in activity.

## Resource Availability

## Acknowledgements

This work was supported by National Institutes of Health (5K08NS105938 to N.A.F.), Brain & Behavior Research Foundation (Award 31729 to N.A.F.) and SFARI Bridge to Independence award (604325 to N.A.F.), R2C Epilepsy Award from CureSHANK, and a generous donation from Edward Elliott. We thank the University of Utah High-Throughput Genomics core for snRNA-sequencing services.

## Author Contributions

N.A.F. and H.W. designed the experiments and analysis, N.A.F., P.D., H.W., and D.K. performed all experiments, N.A.F., H.W., P.D., D.K., B.C.C., and A.C. performed data analysis, N.A.F. acquired funding, N.A.F., H.W., and D.K. wrote and revised the manuscript.

## Methods

### Animals

Mice were maintained on a 12-hour light-dark cycle (light on at 06:00, light off at 18:00) with access to food and water ad libitum. The temperature was maintained at 22±1°C and the humidity between 30% and 50%. All mice in this study were adults aged more than 3 months. Mice were group housed until isolation at the time of lens implantation or 3 days prior to transcriptomic experiments. Transcriptomic experiments include 10 C57BL/6J mice (Jax Strain No.000664), 4 Shank3B WT and 5 Shank3B KO (Jax Strain No.017688^36^). Imaging experiments include 10 Shank3B WT and 11 Shank3B KO, half of which are Ai75D (Jax Strain No.025106) heterozygous. Shank3B WT and KO littermates were generated through crosses between Shank3B-heterozygous parents or Shank3B-heterozygous;Ai75D heterozygous parents. Novel juveniles used in the social interaction assays were sex-matched and aged 4-6 weeks. All experiments were conducted according to the National Institutes of Health guidelines, and protocols were approved by the University of Utah Institutional Animal Care and Use Committee.

### Stereotactic Injection and Lens Implantation

Mice were anesthetized with 1.5-2% isoflurane and mounted in a stereotactic frame (Kopf). To image pan-neuronal prefrontal ensemble activity during freely moving behavior, mice were injected with 900 nl of AAV1-syn-GCaMP7f (Addgene, Catalog No.104488-AAV1) unilaterally to the mPFC at anterior-posterior (AP) +1.7, medial-lateral (ML) 0.3, and dorsal-ventral (DV) −2.5 to −2.25 mm from the bregma. To label mPFC neurons innervated by the vHPC, 900 nl of AAV1-hSyn-Cre (Addgene, Catalog No.105553-AAV1) was injected into the vHPC at AP −3.0, ML 3.0, and DV −4.4 to −3.9 mm from the bregma. To label mPFC neurons projecting to the NAc, 900 nl of AAVretro-hSyn-mCherry (Addgene, Catalog No.114472-AAVrg) were injected into the NAc at AP 0.9, ML 0.65, DV −4.8 to −4.6 mm from the bregma.

After at least 1 week of recovery, tissue above the mPFC was aspirated, and a GRIN lens with an integrated baseplate (1.0 × 4.0 mm, Inscopix) was implanted at the same AP and ML coordinates of the mPFC to a depth of 2.1 mm. Imaging experiments were performed at least 4 weeks after lens implantation to permit recovery.

### Social Behavior and EZM Recordings

All experiments were performed during the light cycle of the animals. Mice were accustomed to the room and handling by the experimenter prior to behavior assays with the microendoscope attached. First, the test mouse was left alone in its home cage for 10 minutes (HC1 epoch). Then, a novel, sex-matched juvenile mouse was introduced to the test mouse’s home cage (Social epoch). After 10 minutes of the Social epoch, a juvenile mouse was removed from the test mouse’s home cage for 10 minutes (HC2 epoch). Lastly, the test mouse was moved to the elevated zero maze where the test mouse explored the maze for 30 minutes (EZM epoch). To match the duration of each epoch, only the first 10 minutes of the EZM epoch were used for further analysis.

### Image Acquisition and Segmentation

Images were acquired using the nVue system (Inscopix) at 20 Hz and spatially downsampled by a factor of 4. The data acquisition system was connected to a computer and synced to behavior video acquisition software (StreamPix) with a transistor-transistor logic (TTL) signal. The light power for the green channel (GCaMP signal) was 0.2 mW for image acquisition. To generate a red image we collected a short (∼100 frame) video of red images at the end of each video, in which the red channel (mCherry or tdTomato signal) was adjusted to maximize signal to noise (typically 0.1-0.3 mW).

The raw images were processed using the Inscopix data processing software. First, the images were spatially down-sampled by a factor of 2, bandpass filtered (0.005 to 0.5 pixels), and corrected for motion artifact. Then, neuronal signals were segmented using the Constrained Nonnegative Matrix Factorization for microEndoscopic Data (CNMF-E) algorithm with the following parameters: cellDiameter = 10, gaussianFilterSize = 1, minCorr > 0.90, minPnr = 10, mergeThreshold = 0.7, patchOverlap = 20, patchSize = 80, ringSizeFactor = 1.4. Segmented neurons were carefully inspected by the experimenters, and outliers with invalid regions of interest (ROIs) or noisier signals were excluded from further analysis. To identify neurons labeled with vHPC input of NAc output, a multicolor registration method was used with the thresholds (lower = 0.10, upper = 0.30).

### Quantifying ensemble activity

So that we could quantify the activity of each neuron during different behaviors, we first converted z-scored calcium traces to binarized rasters, using a threshold of 3. Frames in which the neuronal activity exceeded this threshold were considered to be periods in which neurons were active and assigned a value of 1, whereas other frames the neuron were considered to be inactive. For each behavior, we then calculated the proportion of frames in which each neuron was active. This permitted us to calculate the mean activity (fraction of time active) for each neuron during each behavior.

### Quantifying behaviorally modulated neurons

To identify neurons which were significantly modulated in response to behavior, we utilized our previously published method to compare the activity of each neuron during social interaction or EZM exploration to a null distribution generated by circularly shuffling the calcium activity 10,000 times^9^. Neurons were considered to be positively modulated if they exceeded the 90^th^ percentile of shuffled data, and negatively modulated if the mean activity in the real data was less than the 10^th^ percentile of shuffled data. For spatially binned analysis, neurons were binned into quintiles such that the topmost bin corresponds to > 80^th^ percentile and the lowest bin corresponds to < 20^th^ percentile.

### Quantifying co-modulated Neurons

To identify co-modulated neurons, we identified neurons which were significantly modulated (either positively or negatively) in both behaviors. This result was relatively sparse. To identify neurons which increased their activity in response to both behaviors we identified those neurons which were positively modulated (> 50^th^ percentile) during both behaviors.

### Social interaction assay for transcriptomic analysis

One hour prior to euthanasia for nuclei isolation, the social cohort was subject to a social interaction assay in which they were sequentially exposed to 5 novel juveniles in the home cage. Novel juveniles were sex matched, and each interaction lasted for 5 minutes for a total of 25 minutes of social interaction. Control groups were exposed to sequential cage lid openings to mimic the experience of the social cohort. Thirty-five minutes after the final social interaction (or final lid opening), mice were euthanized by isoflurane overdose followed by immediate decapitation and tissue dissection. To reduce IEG expression induced by social interactions with cage mates, all mice were socially isolated for three days prior to this assay. Mice were additionally habituated to the behavior room and lid opening for about 10 seconds every 5 minutes for two hours on the day prior to the social assay. To prevent light-cycle induced effects on IEG expression and neuronal activity, all behavioral experiments were performed between 6:00 and 7:00 AM, and tissue was harvested between 7:00 and 8:00 AM for all animals.

### Single-nuclei RNA sequencing

Mice were deeply anesthetized under 5% isoflurane, negative toe pinch was confirmed, and they were decapitated. Brains were quickly removed and submerged in ice cold 1X phosphate buffered saline with 15 uM actinomycin-D to prevent artificially induced transcriptional perturbation^37^. The mPFC was micro-dissected and nuclei were isolated following our previously described^38^. Briefly, regions were homogenized with a pestle in lysis buffer containing 15 uM actinomycin-D (Sigma-Aldrich, Catalog No. A1410-10MG) and Triton-X (MilliporeSigma, Catalog No. NUC201) and spun down. Supernatant was removed and replaced with new lysis buffer and samples were spun down two more times. Samples were then filtered through a 40 uM Flowmi cell strainer (Fisher Scientific, Catalog No.14100150) and myelin was removed using a sucrose gradient (MilliporeSigma, Catalog No. NUC201). Nuclei were resuspended and spun down in buffer containing 0.2 U/ul RNAse inhibitor and 0.5% Ultrapure BSA (Fisher Scientific, Catalog No.AM2616) to wash off the remaining sucrose. The pellet was finally resuspended, filtered and diluted to 700-1,000 nuclei/ul in fresh resuspension buffer. Nuclei were counted on a Nexcellom cell counter. Samples were maintained on ice throughout this process.

Following nuclei isolation, samples were immediately delivered to the University of Utah High-Throughput Sequencing Core for library preparation. Barcoded snRNA cDNA libraries were generated with the 10X Genomics Chromium Next GEM Single Cell 3’ Gene Expression Library prep v3.1. The 10X protocol was followed with the exception of extending the PCR elongation step from 1 to 3 minutes^39^. The cDNA libraries were then sequenced on the Illumina NovaSeq 6000 and 200-300M reads per sample were generated. Reads were mapped to the GRCm39 genome with the 10X Genomics Cell Ranger software (v4.0.8).

### Cell type clustering

Filtered gene expression matrices generated by Cell Ranger were clustered using Seurat (v4.2.1) in R (v4.3.0) ^40^. Low quality nuclei with fewer than 800 features or greater than 5% mitochondrial genes were removed. Nuclei with greater than 6000 features were removed to eliminate multiplets. We next utilized the top 30 principal components to calculate the shared nearest neighbors at 0.08 resolution, followed by UMAP dimensionality reduction and visualization. Following clustering, 3 wild-type samples were removed as the tissues were collected at an earlier time point post social interaction, which leads to differences in IEG expression^41^. Any clusters that did not appear in a minimum of 75% of samples or in which 75% or more of the cells belonged to a single animal were removed. Clusters were assigned to known cell types using cell-type-specific marker genes.

### Quantification of a distributed social ensemble

Active neurons were identified in social and control conditions using our previously described criteria for IEG+ cells in the mPFC^41^. IEG+ neurons were identified as those expressing greater than three IEGs (from the panel of 43 IEGs) above the 95^th^ percentile of expression in the wild-type non-social control. The proportion of each identified cell type that met this criterion was calculated by dividing the number of IEG+ cells in a given cluster by the total number of cells in that cluster for each condition (Non-social WT, Social WT, Non-social KO, and Social KO). The fold-increase in the social condition was then calculated by dividing the IEG+ proportion in the social group by the non-social group of the same genotype. The difference between the change in activity induced by social interaction in KO mice as compared to WT was then calculated by dividing the fold change in KO social by WT social.

### Quantification and statistical analysis

Mean calcium activity was calculated for each neuron during each behavior and compared across mice using 2-way ANOVA, as specified. Differences between the social and control group IEG positive proportions for each genotype were determined by comparing the real proportional increase between experimental and control groups to a distribution of shuffled values for each cell type. For shuffling, the number of real cells belonging to each experimental group was selected from within each cell type, and the proportion of IEG+ cells in each selected cell population was calculated. The same comparisons were then calculated for the shuffled data as was done in real data. This was repeated 1000 times to create a distribution of 1000 shuffled values for each cell type. A real value above the 95% of shuffled was considered significant. The significance of the proportional change in the number of active cells in the Shank3 KO was calculated by dividing the shuffled distribution of KO by WT and determining the proportion of the shuffled values the real value was greater than. Cell type proportions were calculated in Prism (v10.4.0) using an ordinary two-way ANOVA with Sidak’s multiple comparisons.

---

**Supplementary Figure 1:**
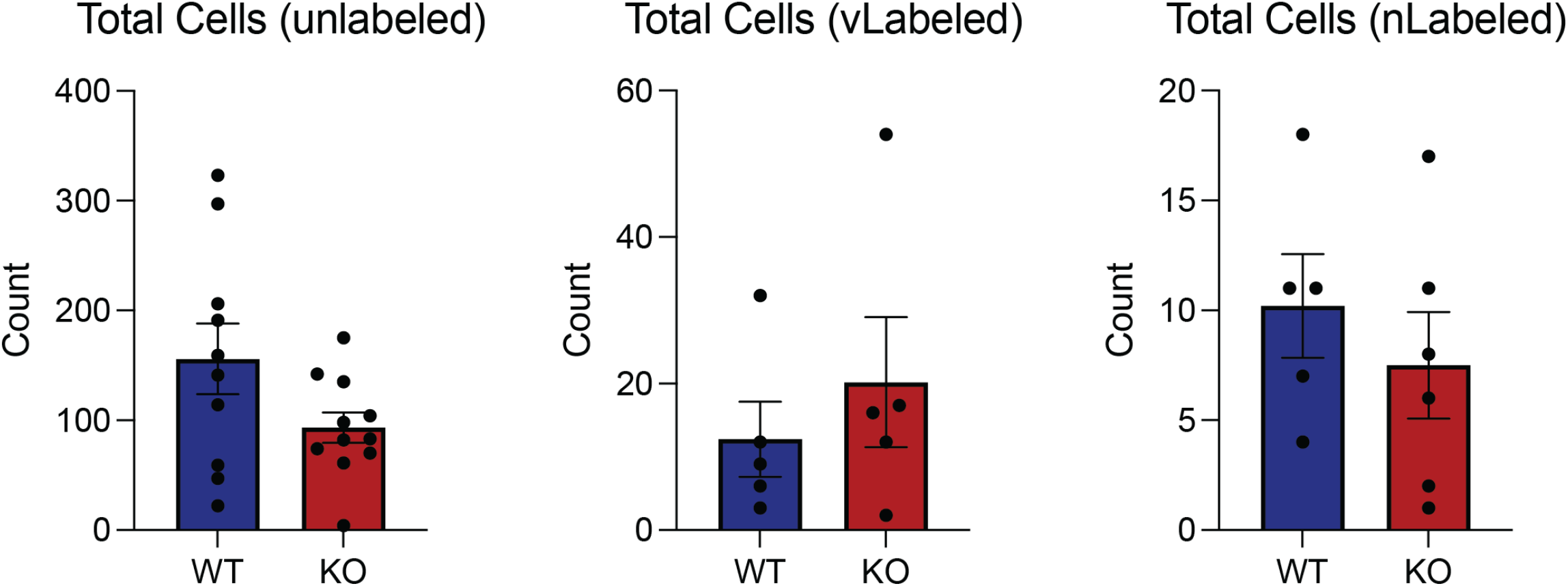
Quantification of Labeled and Unlabeled Cell Populations.

**Supplementary Figure 2:**
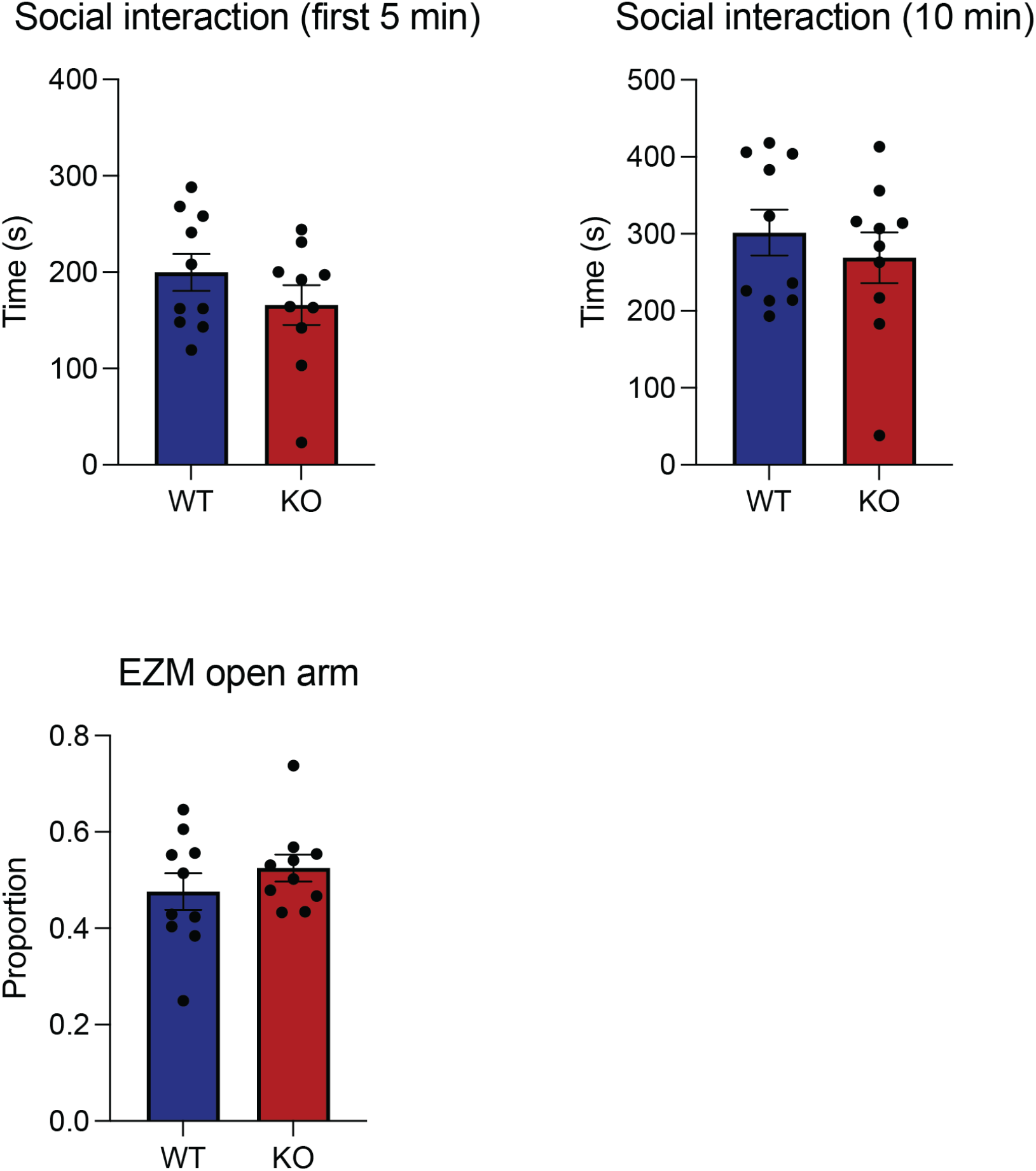
Quantification of Behavior Data.

**Supplementary Figure 3:**
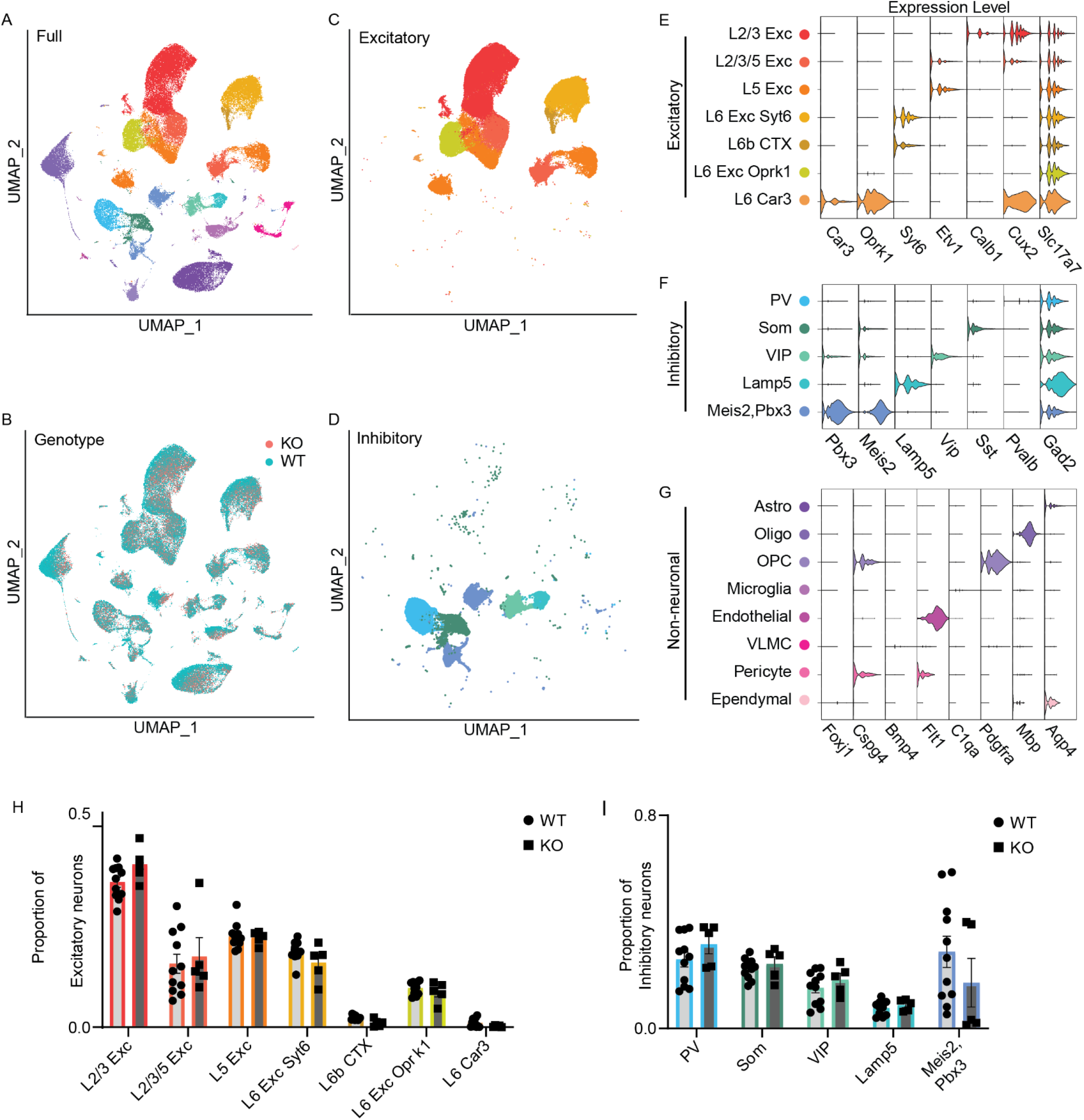
Identification of excitatory and inhibitory neuron subtypes. (A&B) UMAP of identified cell types (n = 116,791 cells; 11 WT and 5 KO mice). (C&D) Excitatory and inhibitory cell clusters. (E) Gene marker expression for each excitatory neuron subtype. (F) Gene marker expression for each inhibitory neuron subtype. (G) Gene marker expression for each non-neuronal subtype. (H) Proportion of cells belonging to each excitatory subtype for WT (light grey) and KO (dark grey) mice (mean +/−SEM). (I) Same as H but for inhibitory neuron subtypes.

**Supplementary Figure 4:**
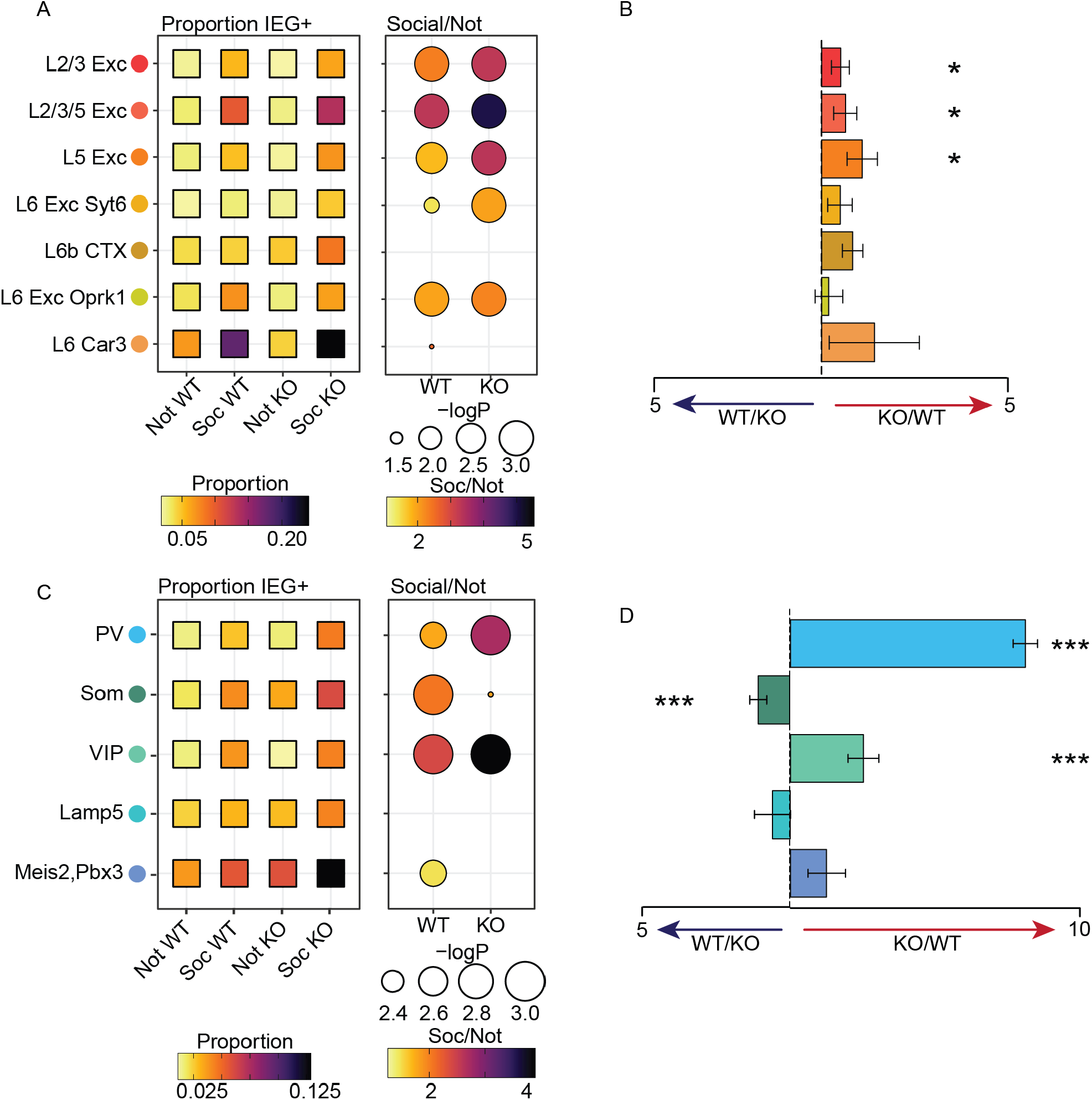
Ensemble recruitment in heterogeneous cell types is altered in Shank3 KO mice. (A) Boxes on the left are colored by the proportion of IEG+ cells for each genotype and cell type before and after social interaction. Circles show the ratio of IEG+ cells in the social condition over the baseline level for each genotype with darker colors showing a greater increase in proportion of active cells. (B) Magnitude of IEG positivity in Shank3 KO animals as compared to WT controls. Bar graph of the ratio of the fold-difference in the proportion of IEG+ neurons in each cell population plotted as WT/KO (Left side) or KO/WT (Right side) to show the increased IEG+ proportion in either KO or WT. (C&D) Same as A&B but for inhibitory neuron populations. P-values and standard deviations calculated by from a null distribution generated by shuffling real data 1,000 times; n = 55,183 WT neurons from 11 WT mice, 29,592 KO neurons from 5 KO mice.

**Supplementary Figure 5:**
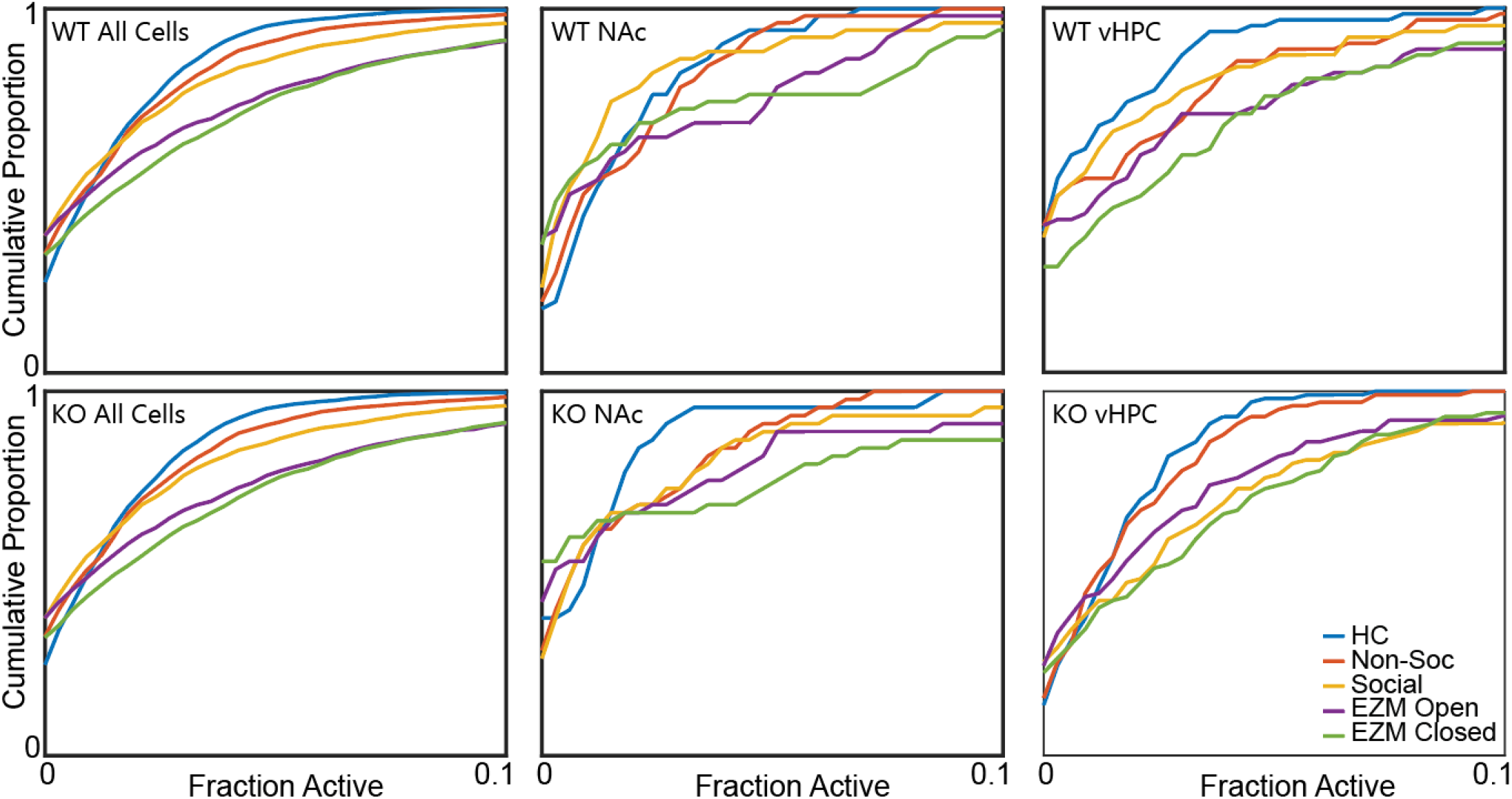
mPFC activity is behavior-dependent. CDF plots of activity for all WT (top, left) and KO (bottom, left) cells, NAc projections from WT (top, middle) and KO (bottom, middle), and vHPC-innervated neurons from WT (top, right) and KO (bottom, right) mice. Data is from 1559 unlabeled neurons from 10 WT mice, 62 vHPC innervated neurons from 5 WT mice, and 51 NAc projecting neurons from 5 WT mice, 1028 unlabeled neurons from 11 KO mice, 101 vHPC innervated neurons from 5 KO mice, and 45 NAc projecting neurons from 6 KO mice.

